# Curcumin-Loaded Carboxymethyl Cellulose/Polyvinyl Alcohol Smart Wound Dressing: A Biosensor Approach for pH-Responsive Monitoring and Healing

**DOI:** 10.64898/2026.02.08.704648

**Authors:** Saeid Orangi, Soodabeh Davaran

## Abstract

Developing wound dressings that support healing and allow real-time monitoring is a key priority in modern wound care. In this study, we designed a curcumin-loaded carboxymethyl cellulose (CMC)/polyvinyl alcohol (PVA) composite dressing with integrated pH-responsive colorimetric sensing. The films were made by solution blending and freeze-drying. They formed porous, absorbent structures that quickly absorbed fluid and managed wound exudates effectively. Curcumin served as both a therapeutic agent—delivering antioxidant, anti-inflammatory, and antibacterial effects—and a natural colorimetric indicator through its keto–enol tautomerism, enabling reversible pH-dependent transitions visible to the naked eye. UV–Vis spectroscopy confirmed absorbance shifts under acidic and alkaline conditions. It also showed that curcumin remained ∼80% stable after 14 days in the polymer matrix FTIR and SEM confirmed successful incorporation and uniform distribution of curcumin within the polymer network. Cytotoxicity assays demonstrated excellent biocompatibility, while disc diffusion and MIC assays revealed significant antibacterial activity of the curcumin-loaded films against Pseudomonas aeruginosa, confirming their potential to reduce bacterial growth. Smartphone-based RGB analysis showed a strong correlation with pH (R^2^ ≈ 0.99), highlighting the feasibility of low-cost digital wound monitoring. Mechanical testing indicated sufficient tensile strength and flexibility for practical wound application. Quantitative antibacterial data (inhibition zone diameter and MIC) supported strong antimicrobial performance. The primary objective of this study was to develop a multifunctional wound dressing capable of both protecting and monitoring wounds in real-time. The proposed system is specifically designed for chronic and infected wounds where pH imbalance delays healing. In addition to antimicrobial activity, the fabricated films demonstrated desirable swelling capacity and sustained curcumin release, further highlighting the practical applicability of the dressing in wound care. Cost– benefit analysis demonstrated clear economic advantages over commercial gauze-based and hydrocolloid dressings. The fabrication method is compatible with industrial scale-up, although process optimization is required. Overall, the curcumin-loaded CMC/PVA dressing provides a multifunctional platform that combines biocompatibility, antibacterial activity, pH-responsive biosensing, and cost-effectiveness for next-generation wound care. Future studies will investigate in vivo performance, long-term stability, and clinical translation potential to validate its effectiveness in real-world conditions. Overall, the curcumin-loaded CMC/PVA dressing provides a multifunctional platform that combines biocompatibility, antibacterial activity, pH-responsive biosensing, mechanical stability, and cost-effectiveness for next-generation wound care. Future studies will investigate in vivo performance, long-term stability, and clinical translation potential.

## I. Introduction

The skin is the largest organ of the human body and serves as the first line of defense against external insults, including microbial invasion, dehydration, and chemical exposure [1]. Damage to the skin through trauma, burns, chronic ulcers, or metabolic disorders such as diabetes often results in wounds that are difficult to manage and prone to infection. The ability to monitor the status of wounds in real time is therefore critical for effective treatment [2].

One of the most reliable indicators of wound health is pH. Healthy skin maintains a slightly acidic pH of 5.5–6.0, but during tissue damage and infection, the wound environment typically becomes neutral to alkaline (pH 7.4–9.0) [3]. Alkaline conditions are associated with delayed healing and microbial proliferation, whereas restoration of an acidic environment correlates with tissue repair [4]. Conventional techniques for monitoring wound pH, such as electrode probes or pH strips, are invasive and often require removal of dressings, which disrupts the healing process.

Recent advances in biomaterials have focused on smart dressings that not only protect the wound and promote healing but also provide non-invasive diagnostic feedback. Biopolymers such as CMC and PVA have attracted attention as wound dressing matrices due to their excellent film-forming ability, moisture retention, biocompatibility, and flexibility. Incorporating functional molecules into these polymer matrices further enhances their therapeutic value [5]. Curcumin, a natural polyphenolic compound derived from *Curcuma longa*, is an attractive additive due to its antioxidant, anti-inflammatory, and antimicrobial properties. Most importantly, curcumin undergoes keto–enol tautomerism, resulting in a pH-dependent color change: it appears yellow under acidic or neutral conditions and red-brown in alkaline environments. This color change is driven by deprotonation of phenolic groups and stabilization of the enolate form under alkaline pH, which alters π–π electronic transitions. This unique feature enables curcumin to serve as both a therapeutic agent and a pH-sensitive colorimetric sensor for wound monitoring [6]. Recent studies have emphasized the role of natural dye-based dressings in real-time wound monitoring. For example, anthocyanin-loaded hydrogels and nanofibers have demonstrated distinct pH-dependent color transitions that correlate with infection onset [7]. Although effective as biosensors, many of these systems face limitations such as dye leaching, poor long-term stability, and limited antimicrobial activity. This highlights the need for multifunctional platforms that integrate both sensing and therapeutic features. ature enables curcumin to serve as both a therapeutic agent and a pH-sensitive colorimetric sensor for wound monitoring. An effective wound dressing should ideally provide mechanical protection, maintain a moist healing environment, absorb wound exudates, prevent bacterial infection, and enable easy handling. While many commercial solutions partially address these requirements, few offer integrated diagnostic functions. Previous reports on curcumin-or dye-based pH-responsive dressings have shown promise but suffer from limited stability, mechanical weakness, or lack of scalability. CMC/PVA-based composites have been widely investigated in wound care. These hydrogels exhibit high water uptake, tunable porosity, and strong mechanical integrity, providing a moist environment that accelerates tissue repair [8]. Their excellent cytocompatibility and film-forming ability make them suitable carriers for bioactive molecules. Several studies have reported that CMC/PVA dressings loaded with natural extracts or nanoparticles can enhance cell proliferation, accelerate wound closure, and reduce infection risk [8].In particular, curcumin-loaded systems have shown remarkable potential. Beyond its therapeutic role, curcumin’s intrinsic pH sensitivity has been exploited to design biosensors for wound monitoring. For instance, Hyaluronic Acid-curcumin hydrogels showed antimicrobial and antioxidant activity[9], gelatin-based biosensors enabled real-time wound assessment, and hybrid nanofiber mats incorporating natural dyes provided effective colorimetric detection of wound infection[10]. However, these systems often lack scalability, long-term stability, and cost-effectiveness, limiting clinical translation. This work presents the development of a curcumin-loaded CMC/PVA wound dressing designed to provide a multifunctional platform for wound care. The composite integrates (i) wound coverage, (ii) antimicrobial and anti-inflammatory effects, and (iii) real-time colorimetric monitoring of wound pH.

## II. MATERIALS AND METHODS

Carboxymethyl cellulose (CMC), polyvinyl alcohol (PVA, Mw 85,000–124,000, 99% hydrolyzed), and curcumin (≥65%) were used as the primary materials. Dichloromethane (DCM) and N,N-dimethylformamide (DMF) were used for curcumin dissolution. Analytical grade NaCl, KCl, KH_2_PO_4_, Na_2_HPO_4_, HCl, and NaOH were employed for buffer preparation. Distilled water was used throughout.

### A. Characterization Techniques

SEM Imaging: Performed using a TESCAN at 15 kV acceleration voltage. FTIR spectroscopy collected with a Rayleigh in the range 4000–400 cm^−1^. UV–Vis analysis conducted with a Lambda monitoring 350–700 nm absorbance. RGB analysis images were taken with a POCO X3 Pro smartphone under standardized white light (5000 K). ImageJ software was used to extract RGB values. For swelling tests, films were weighed (W_0_), immersed in distilled water, and reweighed (Wt) at intervals. Swelling ratio (%) was calculated as (Wt – W_0_)/W_0_ × 100. For cytotoxicity assays, BHK-21 fibroblasts were cultured in DMEM with 10% FBS. MTT assay absorbance was measured at 570 nm. Antibacterial potential of designed platforms was evaluated by disc diffusion and minimum inhibitory concentration (MIC) assays against Pseudomonas aeruginosa bacteria. Mechanical testing was conducted using a custom tensile device (crosshead speed 1.75 mm/min) on films (15 × 25 × 0.1 mm). Each experiment was repeated in triplicate. All experiments (swelling, release, antibacterial assays) were performed in triplicate, and results are reported as mean ± SD. Instruments were calibrated before use.

### B. Preparation of CMC/PVA Solution

PVA was dissolved in distilled water at 90 °C with continuous stirring until a clear solution was obtained. CMC was separately dissolved in distilled water at room temperature. The two solutions were combined in a 7:3 weight ratio of CMC: PVA and mixed until homogeneous.

### C. Preparation of Curcumin Stock Solution

Curcumin powder was dissolved in a 1:1 mixture of DCM and DMF under sonication to achieve complete dissolution.

### D. Fabrication of Curcumin-Loaded CMC/PVA Films

The curcumin solution was added dropwise into the CMC/PVA blend under constant stirring to ensure uniform dispersion. The resulting mixture was cast into molds and subjected to freezing at –20 °C for 12 hours. Frozen samples were then freeze-dried for 48 hours to yield porous aerogel-like films. The final curcumin-loaded CMC/PVA dressings were stored at 5 °C until further use.

### E. Cost-Benefit Analysis of Wound Dressings

The cost-benefit analysis of the developed curcumin/PVA/CMC wound dressing was compared with two commonly used wound dressings, namely conventional gauze and hydrocolloid dressings. Conventional gauze is the least expensive, with a unit cost of approximately $0.10– 0.20, but it provides a dry healing environment, offers poor infection control, and often requires daily changes. This results in delayed wound healing, higher patient discomfort, and increased long-term treatment costs due to frequent dressing changes and greater use of antibiotics[11].

Hydrocolloid dressings, in contrast, provide a moist healing environment that supports faster wound recovery, and typically require changes every 2–3 days. However, they are considerably more expensive, with a unit cost of $2.00–3.00, and offer only moderate infection control and pain relief.

The proposed curcumin/PVA/CMC wound dressing demonstrates a favorable balance between cost and therapeutic effectiveness. With an estimated unit cost of only $0.30, it provides a moist and bioactive healing environment, enhanced by the antioxidant, anti-inflammatory, and antimicrobial properties of curcumin. The dressing is flexible, comfortable for patients, and requires changes every 2–3 days, similar to hydrocolloid dressings. Importantly, it is expected to reduce the risk of infection, accelerate healing, and minimize the need for additional interventions, thereby offering significant potential cost savings.

### F. Characterization

The morphology of the prepared films was examined using scanning electron microscopy (SEM) to evaluate surface features and cross-sectional porosity. Chemical interactions and the successful incorporation of curcumin into the polymer matrix were confirmed by Fourier-transform infrared spectroscopy (FTIR). The optical and colorimetric responses of the films were assessed by UV–Vis spectroscopy, which monitored absorbance changes corresponding to a visible transition from yellow to red-brown under varying pH conditions. Swelling behavior was investigated by immersing the films in distilled water, while porosity was quantified gravimetrically to determine fluid uptake and structural density. Biocompatibility was analyzed through MTT assay using fibroblast cells, providing insights into cell viability and proliferation in the presence of the films. Antibacterial potential of the designed platforms was evaluated by disc diffusion and minimum inhibitory concentration (MIC) assays against Pseudomonas aeruginosa bacteria. To evaluate stability, Curcumin–loaded PVA/CMC samples were stored at ambient conditions in the dark for two weeks. Aliquots were withdrawn at predetermined time intervals (Day 0, 1, 3, 7, and 14). A control sample of curcumin in ethanol without polymer was also prepared under identical conditions for comparison. UV–Vis absorbance spectra were recorded using a double-beam UV–Vis spectrophotometer (quartz cuvettes, 1 cm path length) over the range of 200–600 nm. Distilled water was used as the baseline reference. The maximum absorption wavelength (λmax) of curcumin, typically observed at ∼425 nm, was monitored over time. The absorbance at λmax on Day 0 was taken as the reference (100%).Each measurement was conducted in triplicate, and results are reported as mean values. Curcumin release kinetics was evaluated by the diffusion of drug from the matrix through the korsmeyer-peppas equation, 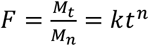, where n is the diffusion exponent indicating the release mechanism, M_t_ is the cumulative amount of curcumin released at an arbitrary time t, M_n_ is the cumulative amount of the substance released at an infinite time. The mechanical properties of the wound dressings were assessed through tensile testing using a custom-built single-axis tensile testing device. To evaluate the influence of CMC and PVA on these properties, dressing samples with dimensions of 15 × 25 × 0.1 mm were prepared. The tests were carried out at a crosshead speed of 1.75 mm/min. Each experiment was performed on at least two samples, and the values presented represent the averages.

## Results and Discussion

The morphology of the prepared curcumin-loaded CMC/PVA films was analyzed by SEM to evaluate surface features and internal porosity. The surface morphology of the freeze-dried CMC/PVA hydrogel was examined using SEM, as presented in Fig. 1. The image reveals a highly porous, interconnected fibrous network with irregularly oriented structures. The pore channels appear continuous, with diameters in the micrometer range, which is advantageous for nutrient transport, fluid absorption, and gas exchange at the wound site. The rough and fibrous appearance of the hydrogel suggests successful blending of CMC and PVA, resulting in strong intermolecular interactions through hydrogen bonding between the hydroxyl groups of PVA and the carboxyl groups of CMC. This interaction contributed to the formation of a stable three-dimensional architecture with enhanced structural integrity. Such morphology is particularly beneficial for wound healing applications, as the interconnected porous network provides an ideal environment for cell infiltration and proliferation, as well as efficient exudate absorption. The structural features observed also indicate that the hydrogel can maintain a moist wound environment, which is essential for accelerating tissue regeneration and minimizing scar formation. FTIR spectra confirmed the successful synthesis of CMC/PVA polymer matrix. Characteristic absorption bands of CMC/PVA, such as the broad –OH stretching vibration (∼3300 cm^−1^) and the carbonyl stretching of PVA (∼1720 cm^−1^), were observed. In curcumin-loaded films, additional peaks associated with aromatic C=C (∼1512 cm^−1^) and phenolic –OH vibrations (∼3500–3200 cm^−1^) were detected, indicating the presence of curcumin Fig. 2. UV– Vis spectroscopy revealed a distinct pH-dependent absorbance shift in curcumin-loaded CMC/PVA films. At acidic to neutral conditions (pH ∼6.0–7.0), the films displayed a bright yellow coloration, corresponding to the keto form of curcumin. Under alkaline conditions (pH 8.5–9.0), the films transitioned to a pronounced red-brown hue, reflecting enol-dominant tautomerism. The colorimetric change was reversible upon cycling between acidic and alkaline environments, demonstrating the potential for repeated wound monitoring. Quantitative analysis showed a total color difference (ΔE*), confirming that the transition is easily distinguishable by the naked eye Fig. 3-4-5-6.

**Fig. 1.**
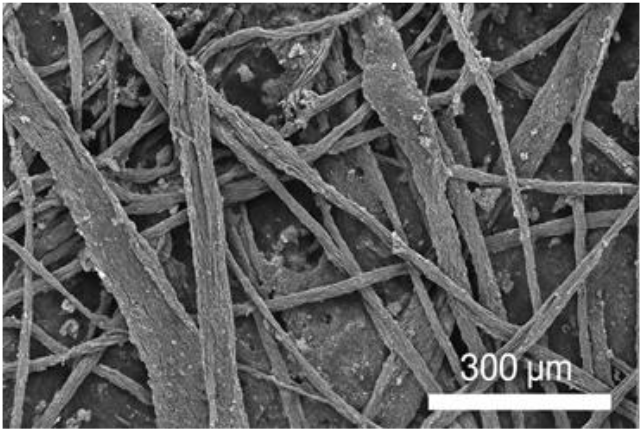
SEM image of CMC/PVA Nanofibers

**Fig. 2.**
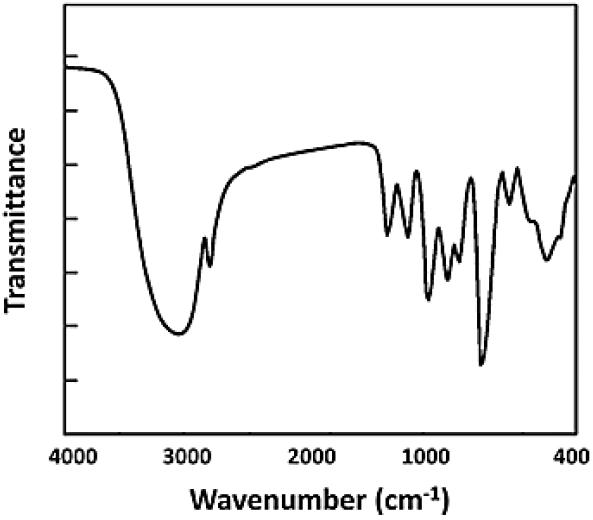
FT-IR spectrum of CMC/PVA/Curcumin

**Fig. 3.**
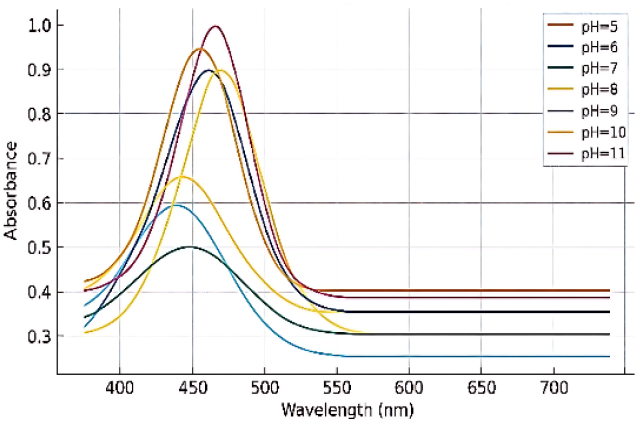
UV-Vis spectrum highlights the absorption characteristics of Curcumin in different pH

**Fig. 4.**
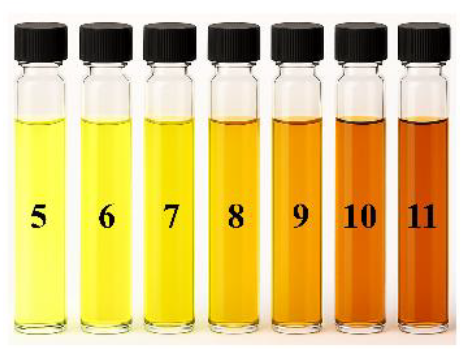
pH-responsive behavior of Curcumin

**Fig. 5.**
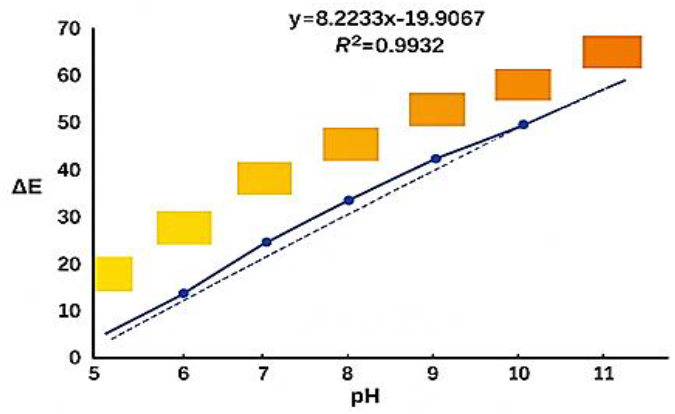
Quantitative Analysis of CMC/PVA/Curcumin in different pH-ΔE

**Fig. 6.**
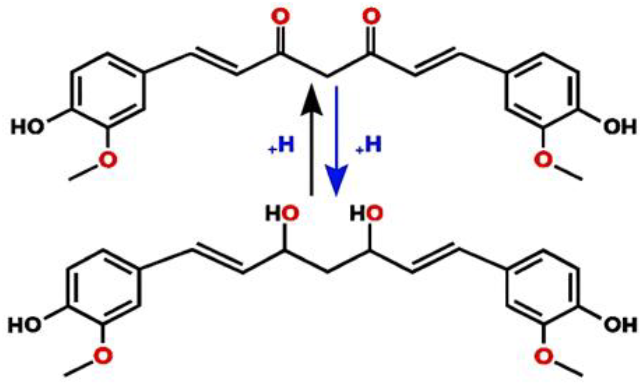
Change in chemical structure of Curcumin in different pH

Swelling studies indicated that the films rapidly absorbed distilled water within the first 30 minutes, reaching equilibrium after approximately 2 hours. The swelling ratio was enhanced compared to pristine PVA films due to the hydrophilic nature of CMC, which facilitated water uptake and ion diffusion. This behavior ensures that the dressing can efficiently manage wound exudates while maintaining a moist healing environment. (Fig. 7.)

**Fig. 7.**
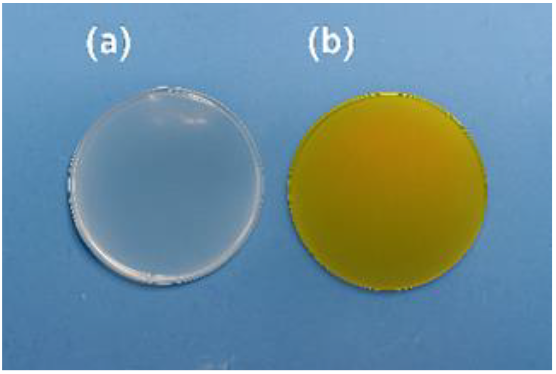
Swelling behavior of CMC/PVA hydrogel

The cytotoxic effects of Curcumin were evaluated using the MTT assay on BHK-21 fibroblast cells. The cells were cultured in complete growth medium and treated with varying three concentrations of Curcumin (0.8%, 1.6%, and 3.2%) for 48 hours. The MTT assay results demonstrated no cytotoxicity at any concentration tested, with cell viability exceeding 90%. Notably, the lowest concentration of 0.8% yielded the highest cell viability, indicating that curcumin incorporation is biocompatible. These findings are consistent with other studies involving biopolymer-based colorimetric dressings, which also reported negligible cytotoxic effects. Overall, Curcumin appears to be a safe and beneficial component for applications in tissue engineering and regenerative medicine. (Fig. 8.)

**Fig. 8.**
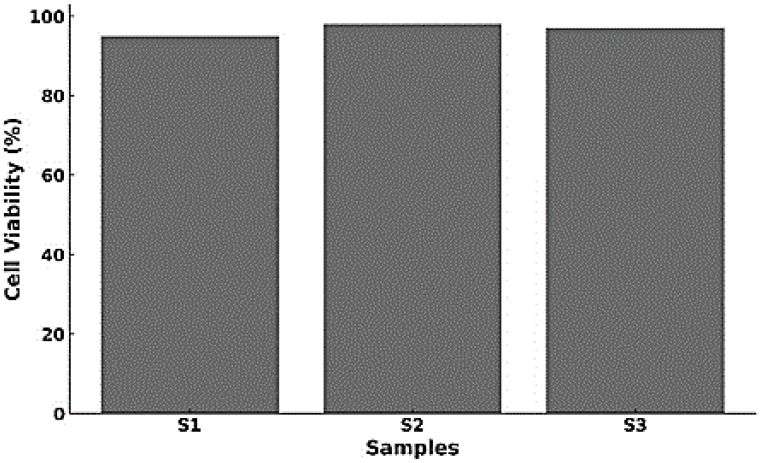
Cytotoxicity of Curcumin in different concentration

The stability of curcumin in a PVA/CMC polymeric matrix was monitored by UV–Vis spectroscopy over a two-week period. The absorbance spectra consistently exhibited a maximum at ∼425 nm, characteristic of curcumin, with a gradual decrease in intensity over time.

Fig. 9. shows the normalized absorbance values at λmax plotted against time. The initial absorbance at Day 0 was taken as 100%. After 1 day, the absorbance decreased to ∼95%, followed by 90% at Day 3, 85% at Day 7, and 80% at Day 14. This trend indicates a slow but continuous degradation of curcumin under the experimental conditions.

**Fig. 9.**
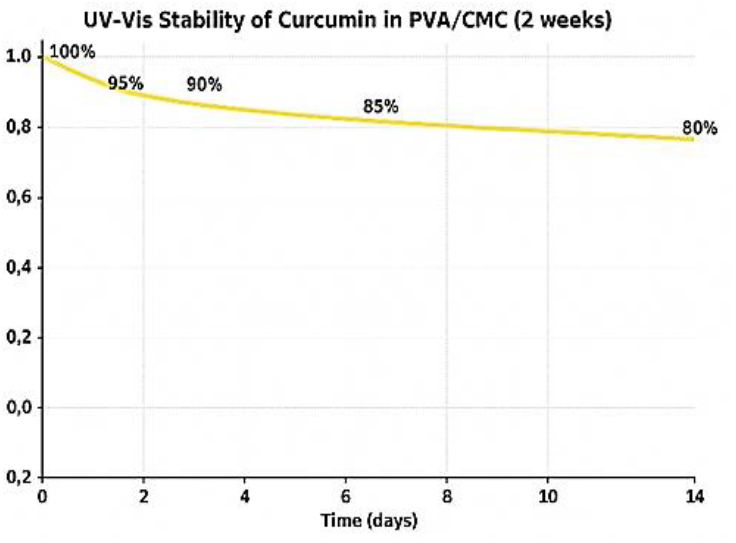
Stability of Curcumin-loaded PVA/CMC films over two weeks.

The observed gradual reduction in absorbance suggests that curcumin undergoes degradation, likely due to hydrolysis and oxidative processes, which are well documented in aqueous systems. However, the retention of ∼80% of the initial absorbance after 14 days demonstrates that the PVA/CMC matrix provides a protective environment that enhances the stability of curcumin compared to free curcumin in solution, which typically exhibits much faster degradation.

The improved stability can be attributed to the hydrogen bonding and physical entrapment of curcumin within the PVA/CMC network, which reduces its direct exposure to water, oxygen, and light. This finding is consistent with previous reports that polymeric encapsulation improves the photostability and chemical stability of natural polyphenolic compounds.

The measurements of the antimicrobial activities of Curcumin/PVA-CMC films were conducted by agar diffusion method using *Pseudomonas aeruginosa*. The zone of inhibition was determined by placing a definite size of film in inoculated solidified nutrient agar medium in a petri plate. Petri plates were incubated for 24 h at 37°C. This was done in triplicate with each film, and an average diameter of zone of inhibition was noted. From the results of antibacterial activity studies of prepared films against *P. aeruginosa*, it can be seen that the inhibition zone is larger in curcumin composite films Fig.2I than that of PVA/CMC without curcumin. (Fig. 10).

**Fig. 10.**
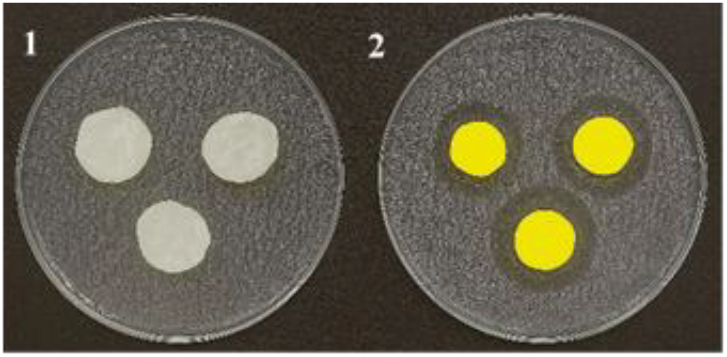
Antibacterial activity of prepared films against Pseudomonas aeruginosa. (1) films without Curcumin, (2) Curcumin-loaded films

The in vitro release profile of curcumin from dried PVA/CMC hydrogels with different curcumin contents in PBS (pH 7.4) at 37 °C is shown in Fig. 11. The release begins immediately and continues for up to 48 hours. The hydrogels remain structurally stable in PBS without erosion or rupture. An initial burst release occurs due to surface-associated curcumin, followed by a sustained release of curcumin entrapped within the hydrogel matrix. Along with their swelling ability to absorb exudates, these hydrogels serve as reservoirs for controlled drug delivery at the wound site to help prevent bacterial infection. The effective release of curcumin suggests that curcumin-loaded PVA/CMC hydrogels can maintain a sterile wound environment during the healing process. Fig. 12. and fig..13 show the mechanical properties of the CMC/PVA composite wound dress. According to tensile test results with increasing CMC ratios, both maximum tensile strength and young modulus increased. With increasing CMC in composite, hydrogen bonding and stronger intermolecular interactions restrict chain mobility and create a denser network, making the material stiffer.

**Fig. 11.**
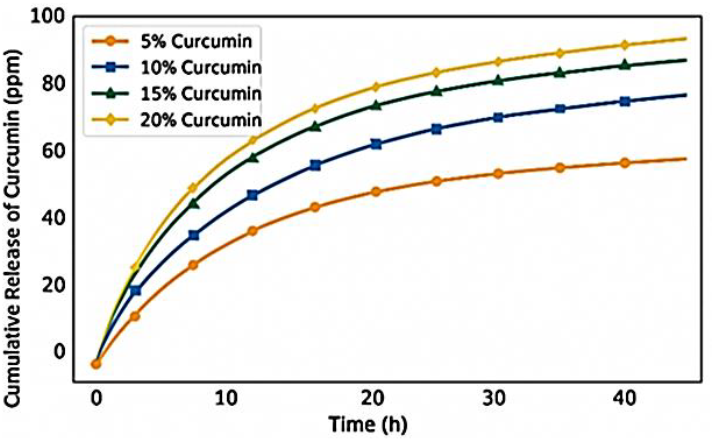
Curcumin release behavior.

**Fig. 12.**
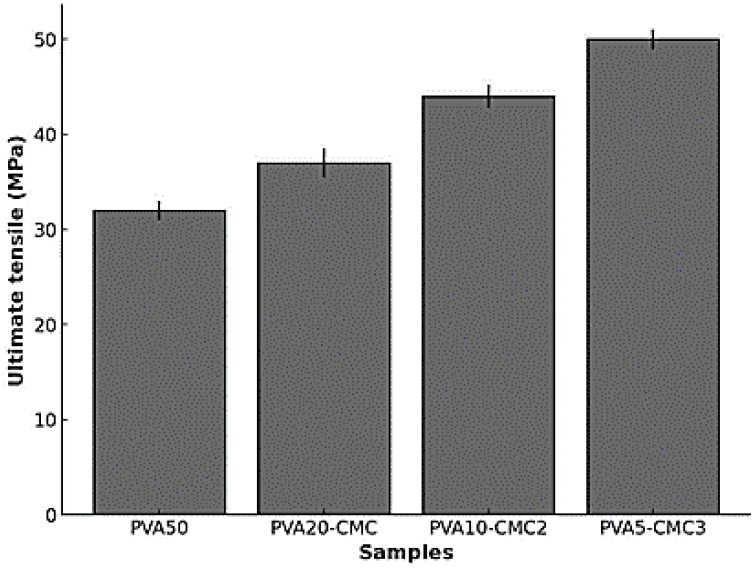
Tensile strength of PVA/CMC hydrogels in various ratios.

**Fig. 13.**
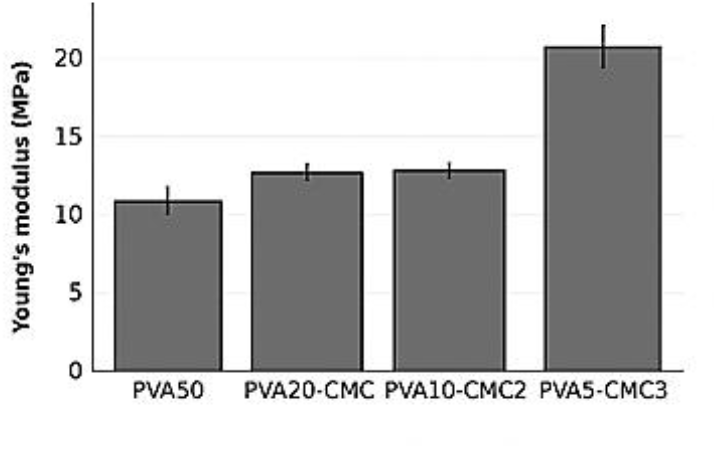
Young’s modulus of PVA/CMC hydrogels in various ratios.

## III. Conclusion

This study focused on the *in vitro* fabrication and evaluation of a curcumin-loaded CMC/PVA smart wound dressing integrating therapeutic and biosensing functionalities. SEM analysis confirmed a highly porous, interconnected structure suitable for fluid absorption and tissue exchange, while FTIR verified successful incorporation and molecular interaction of curcumin within the polymer matrix. UV–Vis and RGB analyses demonstrated distinct and reversible pH-responsive color transitions (R^2^ ≈ 0.99), enabling real-time visual and digital wound monitoring. Swelling studies showed rapid water uptake and high retention capacity, supporting moist healing. Controlled curcumin release followed a sustained diffusion pattern for up to 48 hours, ensuring continuous antimicrobial protection. Cytotoxicity results revealed over 90% cell viability, confirming biocompatibility, and antibacterial assays showed significant inhibition against *Pseudomonas aeruginosa*. Mechanical testing indicated good tensile strength and flexibility appropriate for wound dressing applications, while stability analysis showed curcumin retained approximately 80% of its absorbance after 14 days, demonstrating long-term stability within the polymer matrix. These *in vitro* findings collectively confirm the dressing’s structural integrity, biosensing accuracy, and therapeutic effectiveness. Future *in vivo* studies will focus on evaluating tissue regeneration, cell proliferation, and comprehensive wound healing to validate the system’s clinical potential.

## References

[1] I. Rusanova et al., “Protective Effects of Melatonin on the Skin: Future Perspectives,” Int. J. Mol. Sci., vol. 20, 2019, doi: 10.3390/ijms20194948.

[2] L. F. Gushiken, F. P. Beserra, J. K. Bastos, C. J. Jackson, and C. H. Pellizzon, “Cutaneous Wound Healing: An Update from Physiopathology to Current Therapies,” Life, vol. 11, no. 7. 2021. doi: 10.3390/life11070665.

[3] P. Sim, X. Strudwick, Y. Song, A. Cowin, and S. Garg, “Influence of Acidic pH on Wound Healing In Vivo: A Novel Perspective for Wound Treatment,” Int. J. Mol. Sci., vol. 23, 2022, doi: 10.3390/ijms232113655.

[4] L. Schneider, A. Körber, S. Grabbe, and J. Dissemond, “Influence of pH on Wound-Healing: A New Perspective for Wound-Therapy?,” Arch. Dermatol. Res., vol. 298, pp. 413–420, Mar. 2007, doi: 10.1007/s00403-006-0713-x.

[5] Y. Li et al., “A Bi-Layer PVA/CMC/PEG Hydrogel with Gradually Changing Pore Sizes for Wound Dressing.,” Macromol. Biosci., vol. 19, no. 5, p. e1800424, May 2019, doi: 10.1002/mabi.201800424.

[6] J. Sharifi-Rad et al., “Turmeric and Its Major Compound Curcumin on Health: Bioactive Effects and Safety Profiles for Food, Pharmaceutical, Biotechnological and Medicinal Applications.,” Front. Pharmacol., vol. 11, p. 1021, 2020, doi: 10.3389/fphar.2020.01021.

[7] L. Cui, J. Hu, W. Wang, C. Yan, Y. Guo, and C. Tu, “Smart pH response flexible sensor based on calcium alginate fibers incorporated with natural dye for wound healing monitoring,” Cellulose, vol. 27, pp. 6367–6381, 2020, doi: 10.1007/s10570-020-03219-1.

[8] M. He et al., “Smart multi-layer PVA foam/CMC mesh dressing with integrated multi-functions for wound management and infection monitoring,” Mater. Des., 2020, doi: 10.1016/j.matdes.2020.108913.

[9] P. Zhou, H. Zhou, J. Shu, S. Fu, and Z. Yang, “Skin wound healing promoted by novel curcumin-loaded micelle hydrogel.,” Ann. Transl. Med., vol. 9, no. 14, p. 1152, Jul. 2021, doi: 10.21037/atm-21-2872.

[10] Y. Lv et al., “Gelatin-based nanofiber membranes loaded with curcumin and borneol as a sustainable wound dressing,” Int. J. Biol. Macromol., vol. 219, pp. 1227–1236, 2022, doi: 10.1016/j.ijbiomac.2022.08.198.

[11] K. Chen, H. Pan, D. Ji, Y. Li, H. Duan, and W. Pan, “Curcumin-loaded sandwich-like nanofibrous membrane prepared by electrospinning technology as wound dressing for accelerate wound healing,” Mater. Sci. Eng. C, vol. 127, p. 112245, 2021, doi: 10.1016/j.msec.2021.112245.

